# Incorporating phylogenetic information in microbiome abundance studies has no effect on detection power and FDR control

**DOI:** 10.1101/2020.01.31.928309

**Authors:** Antoine Bichat, Jonathan Plassais, Christophe Ambroise, Mahendra Mariadassou

## Abstract

We consider the problem of incorporating evolutionary information (e.g. taxonomic or phylogenic trees) in the context of metagenomics differential analysis. Recent results published in the literature propose different ways to leverage the tree structure to increase the detection rate of differentially abundant taxa. Here, we propose instead to use a different hierachical structure, in the form of a correlation-based tree, as it may capture the structure of the data better than the phylogeny. We first show that the correlation tree and the phylogeny are significantly different before turning to the impact of tree choice on detection rates. Using synthetic data, we show that the tree does have an impact: smoothing p-values according to the phylogeny leads to equal or inferior rates as smoothing according to the correlation tree. However, both trees are outperformed by the classical, non hierachical, Benjamini-Hochberg (BH) procedure in terms of detection rates. Other procedures may use the hierachical structure with profit but do not control the False Discovery Rate (FDR) *a priori* and remain inferior to a classical Benjamini-Hochberg procedure with the same nominal FDR. On real datasets, no hierarchical procedure had significantly higher detection rate that BH. Although intuition advocates the use of a hierachical structure, be it the phylogeny or the correlation tree, to increase the detection rate in microbiome studies, current hierachical procedures are still inferior to non hierachical ones and effective procedures remain to be invented.

## 1 Introduction

The microbiota, loosely defined as the collection of microbes that inhabit a given environment, has become an increasingly important research topic in the last two decades as it proves to either play an active role or be associated with health conditions (Lynch and Pedersen, 2016; Opstelten et al., 2016). For instance, specific changes in microbiome composition have been associated to Inflammatory Bowel Diseases (IBD) (Morgan et al., 2012) and liver cirrhosis (Qin et al., 2014). The microbiota also influences efficiency of cancer therapy (Routy et al., 2018) and there is a growing interest in finding biomarker microbes that could be used predict the response to treatment (Behrouzi et al., 2019). The effect of the microbiota is not limited to human health: works in plant biology show that the root microbiota can improve resistance to stress (Trivedi et al., 2017). Molecules produced by the microbiota can also have a profound impact on stress tolerance (Bernardo et al., 2017), plant health (Mendes et al., 2011) and pathogen control (Bartoli et al., 2018).

There are two main approaches to profile the microbiome using sequence data: amplicon sequencing and whole genome shotgun (WGS) sequencing. In amplicon sequencing, a marker-gene that acts as a “barcode” (*e.g.* the 16S rRNA gene) and carries taxonomic information about the bacteria is first amplified and then sequenced whereas in WGS sequencing, the whole metagenome is sequenced with no prior amplification of a specific region. Although WGS sequencing is less affected by technical bias than amplicon sequencing and can profile both taxonomic and functional composition of the microbiome, it suffers from higher costs and requires complex bioinformatics pipelines. We focus in this work on taxonomic profiles.

In the amplicon approach, sequence reads are first clustered into Operational Taxonomic Units (OTUs) using either a 97% sequence similarity threshold (Caporaso et al., 2010), threshold-free agglomerative approaches (Mahé et al., 2015; Escudié et al., 2017) or divisive approaches to produce taxonomic oligotypes (Eren et al., 2015) or Amplicon Sequence Variants (ASVs) (Callahan et al., 2016). Divisive and threshold-free agglomerative approaches achieve finer taxonomic resolutions than the threshold-based similarity approach. Using WGS in the ecosystems where a bacterial gene catalog is available, such as the human gut (Li et al., 2014) or the pig gut (Xiao et al., 2016), the standard approach consists in mapping the reads against the catalog and then clustering the bacterial genes based on their abundance profiles to produce metagenomic species (MGS) (Nielsen et al., 2014) or clusters of co-abundant genes to reconstruct microbial pan-genomes (MSP) (Plaza Oñate et al., 2018). We will refer to taxa, noting that the term can designate OTUs, ASVs, oligotypes, MGSs, MSPs and generally any feature found in abundance tables.

The microbial taxa share a common evolutionary history that can be encoded by a phylogenetic tree. For amplicon sequencing, the phylogenetic tree of taxa can even be reconstructed based on the sequence divergence of taxa (Price et al., 2010). Related taxa are generally thought to perform similar biological functions. For example, Philippot et al. (2010) shows a strong association between taxonomic lineage and ecological niche in soil microbiota. Chaillou et al. (2015) reports similar associations in food microbial ecosystems. This observation prompts the development of several tree-based hierarchical methods, build under the assumption that taxa associated to a phenotype of interest are clustered in the tree (Martiny et al., 2015). Carroll et al. (2014) considers group-based procedures, with groups defined as clades of the tree. Sankaran and Holmes (2014) proposes an implementation of the hierarchical testing procedure of Yekutieli (2008) aimed at leveraging the phylogenetic tree of the taxa to increase statistical power while controlling the False Discovery Rate (FDR). The FDR is unfortunately only known *a posteriori*, and the implemented testing-procedure is limited to one-way ANOVA with no correction for differences in sequencing depths. Matsen IV and Evans (2013) and Washburne et al. (2017) develop phylogenetic eigenvalues decomposition of species compositions for exploratory data analysis. Finally Xiao et al. (2017) uses the tree as a regularization structure to shrink the test statistics of close-by taxa towards the same value. They use a permutational procedure to control the FDR and report good empirical control of the FDR but the method lacks theoretical grounding.

Unfortunately for phylogeny-based methods, the association between ecological niche and taxonomy reported in Philippot et al. (2010) holds for high-rank taxa but breaks down for lower-rank taxa. Furthermore, in a given ecological niche, it is unclear whether the genetic basis of a given phenotype lies in the core genome, shared by many taxa of a phylogenetic clade, or in mobile elements driving adaptation (Kazazian, 2004), and hence more spread out in the tree (Brito et al., 2016). We question in this work the premise that the phylogenetic (or taxonomic) tree is the relevant hierarchical structure to incorporate in differential studies. We argue that the correlation tree, created from co-abundance data, is a better proxy of biological functions and can increase statistical power with no loss of FDR control in comparison to the phylogeny.

Using several metrics (Billera et al., 2001; Robinson and Foulds, 1981) in the treespace and datasets from previous studies (Ravel et al., 2011; Zeller et al., 2014; Chaillou et al., 2015) with both narrow and broad environmental ranges, we study the distance between the phylogenetic tree and the correlation trees. We compare those distances to the average distance between (i) a focal tree (phylogeny or correlation) and a random tree and (ii) between two random trees to investigate the relationship between proximity in the tree and correlated abundances. We then assess the impact of tree selection on differential studies using both extensive simulation studies and reanalysis of previously published datasets. We compare the results obtained with the phylogeny, the correlation tree, and the standard Benjamini-Hochberg correction. Finally, we discuss the pros and cons of using one or the other in hierarchical procedures and some limitations of our work.

## 2 Material and Methods

### 2.1 Trees

We consider in this study different hierarchical structures, or trees: the phylogenetic tree, the taxonomic tree and the correlation tree.

#### Phylogenetic tree

The phylogeny encodes the common evolutionary history of the taxa. In the amplicon context, it is usually reconstructed based on the sequence divergence of the marker-gene (Price et al., 2010) and branch lengths correspond to the expected number of substitutions per nucleotide.

#### Taxonomic tree

When the phylogeny is not avalaible but taxonomic annotations are, we fall back on the taxonomic tree instead. Inner nodes correspond to coarse taxonomic ranks (*e.g.* phylum, class, order, etc). The hierarchical structure is reconstructed from lineages extracted from regularly updated databases like the one from NCBI (Geer et al., 2009). Branch lengths correspond to the number of levels in the hierarchy: *e.g.* a branch between species-level and genus-level nodes has length 1, a branch between species-level and genus-level nodes has length 2. Unlike phylogenetic trees, taxonomic trees are highly polytomic.

#### Correlation tree

The correlation tree is based on the abundance profiles of taxa across samples and built in the following way. We first compute the pairwise correlation matrix, using the Spearman correlation and excluding “shared zeros”, *i.e.* samples where both taxa are absent. We then change this correlation matrix into a dissimilarity matrix using the transformation *x* ⟼ 1−*x*. Finally, we use hierarchical clustering with Ward linkage on this matrix to create the correlation tree. Branch lengths correspond to the dissimilarity cost of merging two subtrees.

### 2.2 Distances between trees

We consider two different distances between trees: the Robinson-Foulds distance, or RF (Robinson and Foulds, 1981), the Billera-Holmes-Vogtmann distance, or BHV, (Billera et al., 2001). Those distances are computed using different characteristics of the tree (topology, branch lengths, etc) and emphasize different features.

The RF distance is defined on topologies, *i.e.* trees without branch lengths, and based on elementary operations: branch contraction and branch expansion. A branch contraction step creates a polytomy in the tree by shrinking a branch and merging its two ending nodes whereas a branch expansion step resolves a polytomy by adding a branch to the tree. For any pair of trees, it is possible to turn one tree into the other using only elementary operations. The RF distance is the smallest number of operations required to do so. Note that the RF distance gives the same importance to all branches, no matter how short or long.

The BHV distance is defined on trees and accounts for both topology and branch length. It is based on an embedding of tree into a treespace with a complex geometry. All trees with the same topology are mapped to the same orthant, and hyperplanes share a common boundary if and only if they are at RF-distance 2 (one contraction and one expansion step away). For any pair of trees, there is a path in treespace between those two trees. The BHV distance is the length of the shortest of these paths. It can be thought of as the generalization of the RF-distance that upweights long branches and downweights short branches.

### 2.3 Forest of trees

We generated a forest of boostrapped trees and a forest of random trees in the following way. For the boostrapped forest, we generated *N*_*B*_ bootstrap datasets using resampling with replacement (Felsenstein, 1985; Wilgenbusch et al., 2017). Each bootstrap dataset was used to compute a correlation matrix and a correlation tree as detailed in Sec. 2.1.

Random trees were generated from a seed tree by shuffling the leaves labels. This allowed us to generate a forest of random trees with the same number of branches as the seed tree. This is especially important for RF-distances as they scale with the number of branches and we want to study both non-binary taxonomic trees with a high number of polytomies and low number of branches and binary correlation trees, with a high number of branches. We generated *N*_*T*_ random trees from the taxonomic tree and *N*_*C*_ from the correlation tree.

### 2.4 Testing tree equality

The correlation tree is reconstructed from abundance profiles rather than molecular sequences and/or lineages and may therefore be poorly estimated. We use the bootstrap forest to compute a confidence region around the correlation tree. The random trees were used to create a null distribution of distances between random trees.

The full set of 2 + *N*_*B*_ + *N*_*T*_ + *N*_*C*_ trees was used to construct BHV and RF distance matrices. The distance matrices were then used to visualize a 2D-projection of all trees via Principal Coordinates Analysis (PCoA) (Gower, 1966; Jombart et al., 2017; Wilgenbusch et al., 2017). Bootstrap trees were used to test whether the taxonomy was in the confidence region of the correlation tree whereas random trees were used to test whether the taxonomic and correlation trees were closer to each other than to random trees.

We also compared the distance from the correlation tree to each group of trees using a one-way ANOVA.

### 2.5 Differential abundances studies

The literature abounds in differential analysis methods dedicated to abundance data (Soneson and Delorenzi, 2013). Most of them differ in the normalization and preprocessing steps (Dillies et al., 2013). Count data coming from metagenomic studies are very similar to those found in RNA-Seq studies. The former one may exhibit more zeros entries but the same types of normalizations and statistical models can be used for both types of data.

In this paper, the focus is not on normalization and we used most classical approaches in order to assess the impact of taking into account the data hierarchical structure in the differential abundance testing.

We briefly present two methods for differential abundance testing (DAT) that leverage a tree-like structure: *z*-score smoothing as proposed in Xiao et al. (2017) and hFDR as proposed in Yekutieli (2008).

#### 2.5.1 *z*-scores Smoothing

Given any taxa-wise DAT procedure, *p*-values (*p*_1_, … , *p*_*n*_) are first computed for each taxa (leaves of the tree) and then transformed to *z*-scores using the inverse cumulative distribution function of the standard Gaussian. Similarly, the tree is first transformed into a patristic distance matrix (**D**_*i*,*j*_) and then into a correlation matrix **C**_*ρ*_ = (exp (−2*ρ***D**_*i*,*j*_)) between taxa. The *z*-scores **z**= (*z*_1_, … , *z*_*n*_) are then smoothed using the following hierarchical model:

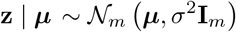

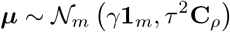

where ***μ*** captures the effect size of each taxa. The maximum a posteriori estimator ***μ***\* of ***μ*** is given by

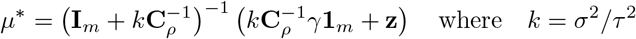

and the FDR is controled using a resampling procedure. This method intuitively pulls effect sizes of taxa close-by in the tree towards the same value. *k* and *ρ* are hyperparameters controling the level of smoothing. Low (resp. high) values of *ρ* (resp. *k*) correspond to high smoothing. Finally, *k*, *γ* and *ρ* are estimated using generalized least-squares.

#### 2.5.2 Hierarchical FDR

Hierarchical FDR (hFDR) considers a different framework where differential abundance can be tested not only for a single taxa but also for groups of taxa, corresponding to inner nodes or clades of the tree. hFDR uses a top-down approach: tests are performed sequentially and only for nodes whose parent node were previously rejected. Formally, the procedure is described in Algorithm 1.

Let ch(*N*) be the children of a node *N*, 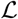 the leaves of the tree, 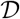 the set of rejected nodes (discoveries), 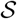 the stack of nodes whose children are yet to be tested and BH_*α*_(*F*) the discoveries within family *F* when testing with a Benjamini-Hochberg procedure at level *α*.

**Algorithm 1.**
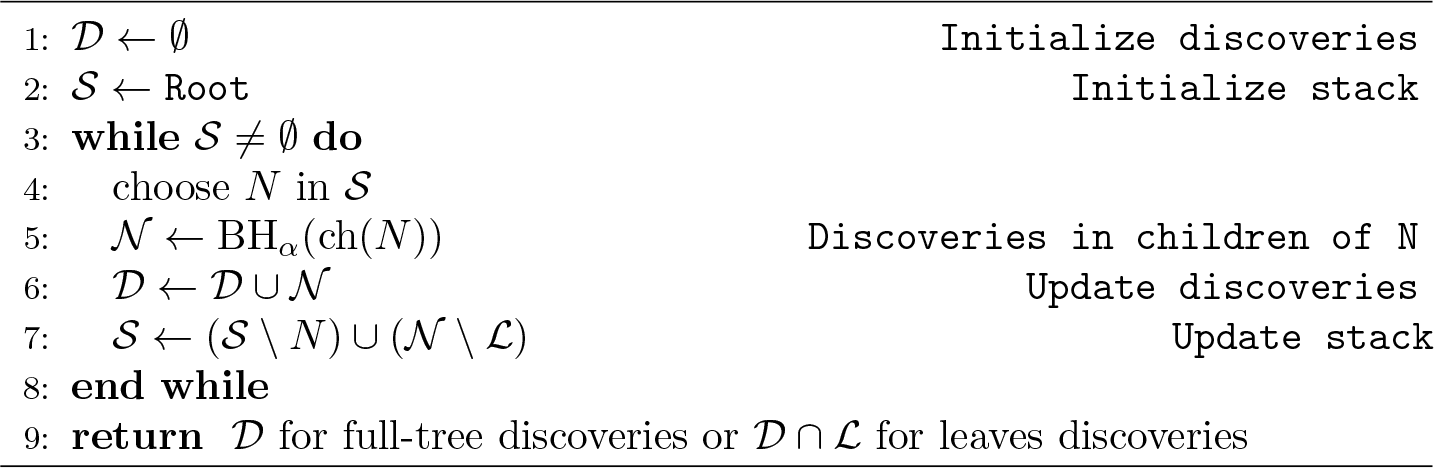
Hierarchical FDR

hFDR guarantees an *a posteriori* global FDR control for leafs at level

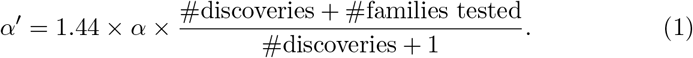

The hFDR procedure is illustrated in Fig. 1.

**Figure 1.**
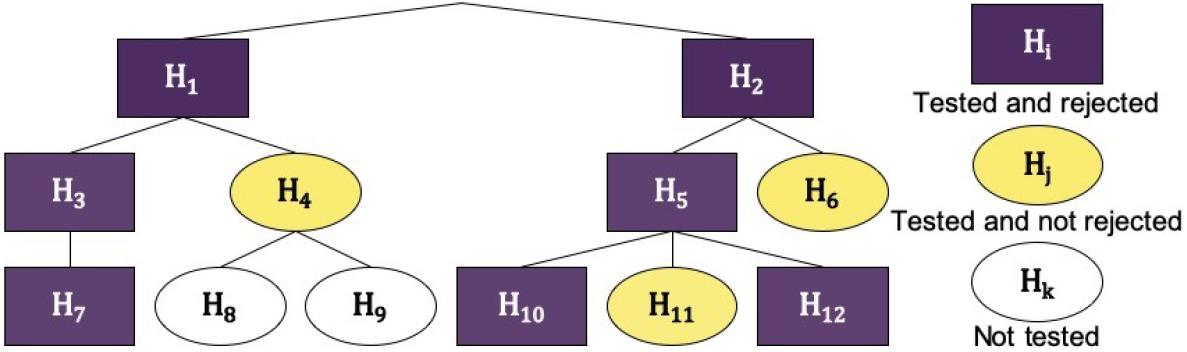
Example wokflow of hFDR. Nodes are numbered from 1 to 12 and the corresponding hypothesis are labeled **H**_1_ to **H**_12_. hFDR first tests and rejects **H**_1_ and **H**_2_. It then tests the family (**H**_3_, **H**_4_), as children of **H**_1_, and rejects **H**_3_ but not **H**_4_. **H**_7_ is tested and rejected, whereas neither **H**_8_ nor **H**_9_ are tested. It proceeds similarly in the tree rooted at node 2. In this example, there are 3 leaf-level discoveries (**H**_7_, **H**_10_ and **H**_12_) and 5 families were tested. Then the *a posteriori* global FDR for leaves is 1.44 × α × 2. Figure adapted from Yekutieli (2008).

#### 2.5.3 Implementations

These two algorithms are implemented in R packages (R Core Team, 2018): `structFDR` (Chen, 2018) for the *z*-scores smoothing and `structSSI` (Sankaran and Holmes, 2014) for hFDR.

The *z*-scores smoothing algorithm as implemented in `structFDR` includes a *fallback* to standard, non hierarchical, independant tests when too few taxa are detected. It was not part of the original algorithm and we therefore used a vanilla implementation, with no fallback (see modified code in `correlationtree` package), to specifically evaluate the impact of the tree in the procedure. `structFDR` requires the user to specify its test. We used non-parametric ones: Wilcoxon rank sum for settings with two groups and Kruskal-Wallis (Hollander and Wolfe, 1973) for settings with three or more groups.

In contrast, the hFDR procedure is only available for one-way ANOVA on the groups, and corresponding *F*-test, and does not correct for differences in sequencing depths. Moreover, we noticed that the global FDR control was off by the corrective factor of 1.44 in Equation (1). We corrected the output of `structSSI` to use the correct FDR values in our analyses.

### 2.6 Methods evaluation

We tested the impact of tree choice on the performance of both procedures (*z*-score smoothing and hFDR) on real data and synthetic data simulated from real dataset in one of two following ways. The code and data used to perform the simulations are available on the github repository github.com/abichat/correlationtree_analysis.

#### 2.6.1 Parametric Simulations

The parametric simulation scheme is based on Xiao et al. (2018). First, a Dirichlet-multinomial model 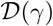 is fitted to the gut microbiome dataset of healthy patients from Wu et al. (2011). Second, a homogenous dataset is created by sampling count vectors *S*_*i*_ from the Dirichlet-Multinomial distribution: (i) a proportion vector *α*_*i*_ is drawn from 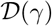, (ii) the sequencing depth *N* is drawn from a negative binomial distribution 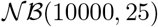 with mean 10000 and size 25 and finally (iii) the counts *S*_*i*_ of sample *i* are sampled from a multinomial distribution 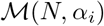.

Differential abundances are then produced as follows. First, each sample is randomly assigned to class *A* or *B*. Second, 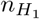 taxa (representing up to 20% of all taxa) were sampled uniformly among all taxa. Finally, the abundances of those taxa are multiplied by a fold-change (chosen in {5, 10, 15, 20}) in group *B*. The process is illustrated in Fig. 2.

**Figure 2.**
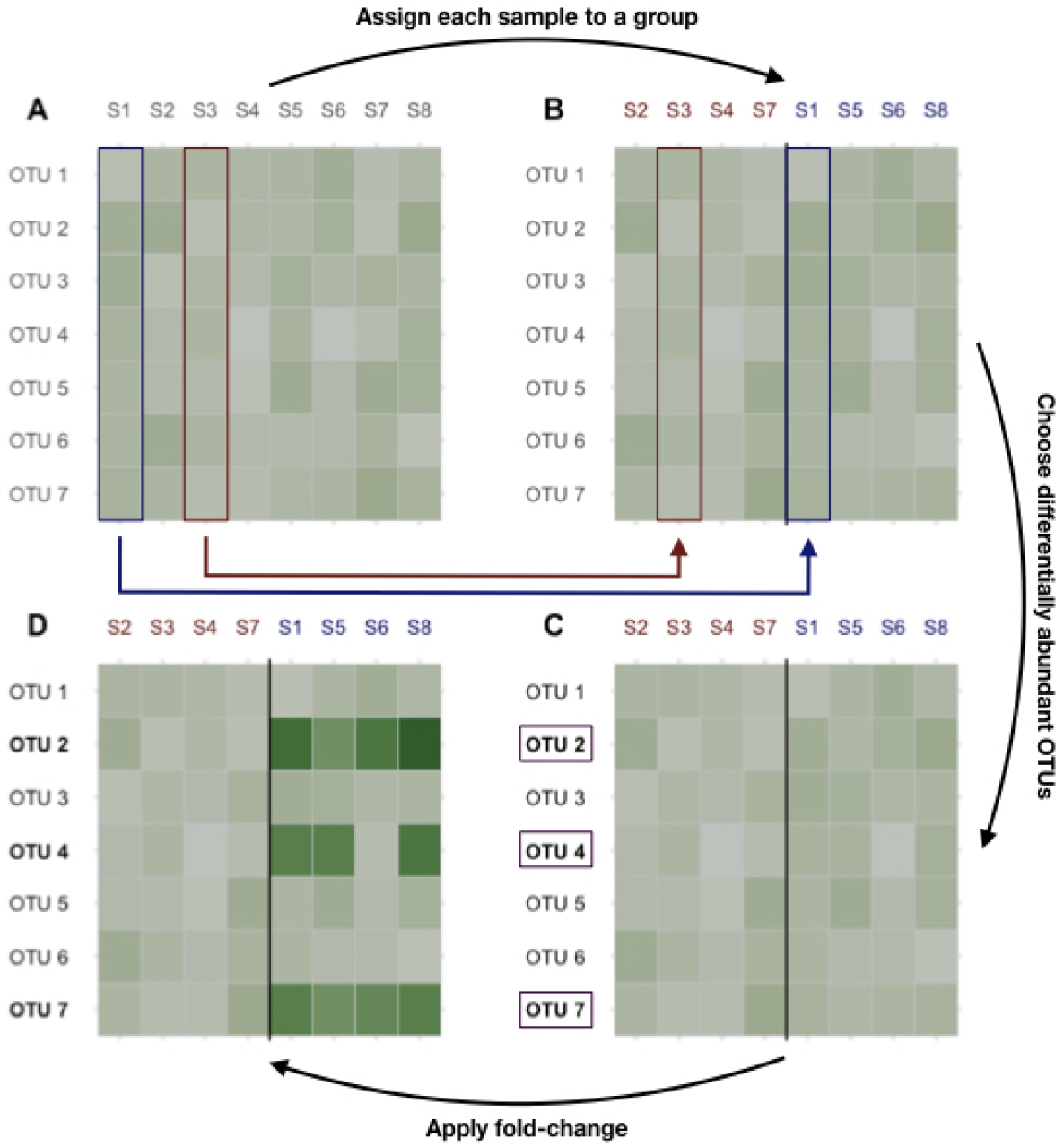
Dataset generation process. A: original count data. B: samples are randomly assigned to class *A* or *B*. C: 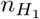 taxa are randomly selected among the most prevalent ones. D: Their abundances are multiplied by the fold-change to produce the final count table.

#### 2.6.2 Non-Parametric Simulations

Non-parametric simulations proceeded like the parametric ones detailed in 2.6.1 with three major differences. First, we used a different dataset with homogeneous samples: the gut microbiome of healthy individuals from North America and Fdji Islands (Brito et al., 2016). Second, we did not fit a Dirichlet-Multinomial to the original dataset but used it as such, to preserve the potential complex correlation structure present in the dataset. Finally, differentially abundant taxa were sampled only from highly prevalent taxa (prevalence ≥ 90%) to ensure that DAT procedures were affected by effect size (fold-change) and hierarchical correction, rather than by sparsity.

#### 2.6.3 Accuracy Evaluation

We used true positive rate (TPR) and FDR to evaluate the performance of *z*-scores smoothing used with five diffent trees: no tree or standard Benjamini-Hochberg (BH), taxonomy, correlation tree, random taxonomy and random correlation tree. BH is our baseline and the random trees are here to evaluate the impact of uninformative trees, with different granularity levels, on the procedure.

We evaluated hFDR by comparing the results obtained using either the taxonomy or the correlation tree in several datasets.

### 2.7 Datasets

We used seven different datasets for the experimental part (see Table 1 for a summary). One was used to study the difference between correlation and phylogenetic trees, one to assess the impact of three choice tree choice on difference abundance testing, three for both and the last two to generate synthetic datasets as described previously. All datasets used in this study are available on the github repository github.com/abichat/correlationtree_analysis.

**Table 1.** Summary table of the different datasets used in this study with information on biome type, taxonomic rank used for the analysis, corresponding number of taxa, number of samples and analyses performed on the dataset: comparison of the correlation and taxonomic trees (Tree), creation of synthetic datasets (Simulations), or impact of the tree on differential abundance procedures (DA).

| Dataset | Biome | Rank | Taxa | Samples | Analysis | Publication |
| --- | --- | --- | --- | --- | --- | --- |
| Chlamydiae | Varied | OTU | 21 | 26 | Tree & DA | Caporaso et al. (2011) |
| Ravel | Vaginal | Genus | 40 | 396 | Tree | Ravel et al. (2011) |
| Wu | Gut | OTU | 400 | 98 | Simulations | Wu et al. (2011) |
| Zeller | Gut | Genus | 119 | 199 | Tree & DA | Zeller et al. (2014) |
| Zeller MSP | Gut | MSP | 878 | 199 | DA | Zeller et al. (2014) |
| Chaillou | Food | OTU | 499/97 | 64 | Tree & DA | Chaillou et al. (2015) |
| Brito | Gut | OTU | 77 | 112 | Simulations | Brito et al. (2016) |

Three of the four datasets used for tree comparison (Ravel, Chaillou and Zeller) were chosen because they are well suited for bootstrapping correlation trees: they had enough samples and enough variability in taxa counts to ensure that a meaningful correlation tree could be computed on bootstrapped datasets. They also represent diverse microbiome with contrasted biodiversity levels: vaginal microbiome for Ravel, food-associated microbiome for Chaillou and gut microbiome for Zeller. Briefly, Ravel et al. (2011) studied a cohort of 396 North-American women from 4 ethnic groups using metabarcoding on the V1-V2 region of 16S rRNA gene. Chaillou et al. (2015) studied food-associated microbiota of 80 processed meat and seafood products using metabarcoding on the V3-V4 region of the 16S rRNA gene. Zeller et al. (2014) considered the gut microbiota of 199 subjects (42 with adenomas, 91 with colorectal cancer and 66 healthy ones), using both shotgun deep sequencing and metabarcoding on the V4 region of 16S rRNA gene. Zeller refers to the 16S rRNA fraction of the data. Details of bioinformatics treatments used to produce abundance count tables are available in the respective publications. All datasets were aggregated at a given taxanomic level and taxa with a prevalence lower than 5% were filtered out.

The fourth one (Chlamidya) was used in Sankaran and Holmes (2014) to assess the performance of hFDR and is an excerpt from the data collected in Caporaso et al. (2011). It consists of bacteria from the Chlamydia phylum and is distributed with `StructSSI` (Sankaran and Holmes, 2014). Finally, the Zeller MSP data originates from the same study as the Zeller data (Zeller et al., 2014). It was created from the shotgun data by reconstructing Metagenomics Species Pan-genomes (MSPs) abundance count table, as reported in Plaza Oñate et al. (2018). Briefly, reads were quality-filtered and unique reads were mapped against the 9.9 million Integrated Gene Catalog (Li et al., 2014) using BBmap (Bushnell, 2014). The gene catalog is organized into 1696 MSPs and each MSPs has set a core genes. The relative abundance of each MSPs was computed by summing the relative abundances of all core genes in that MSP.

The two datasets used to generate synthetic data are the Wu and Brito datasets. The former comes from Wu et al. (2011), a study linking the gut microbiome to alcohol consumption in 98 patients, and was used in Xiao et al. (2017). The latter originates from (Brito et al., 2016), where the gut microbiomes of 81 metropolitan North Americans were compared to those of 172 agrarian Fiji islanders using a combination of single-cell genomics and metagenomics. The metagenomes of Fiji islanders is distributed as part of the R/Bioconductor `CuratedMetagenomicsData` package (R Core Team, 2018; Pasolli et al., 2017) and only the data from the 112 adults were kept, to make it as homogeneous as possible.

## 3 Results and discussion

### 3.1 The Taxonomy Differs from the Correlation Tree

In all studied datasets, the correlation tree is closer to its bootstrap replicates than to either the taxonomy or the randomized trees (Fig. 3, top row). The differences are statistically significant (*p* < 10^−16^, one-way ANOVA with Tukey’s HSD post-hoc test).

**Figure 3.**
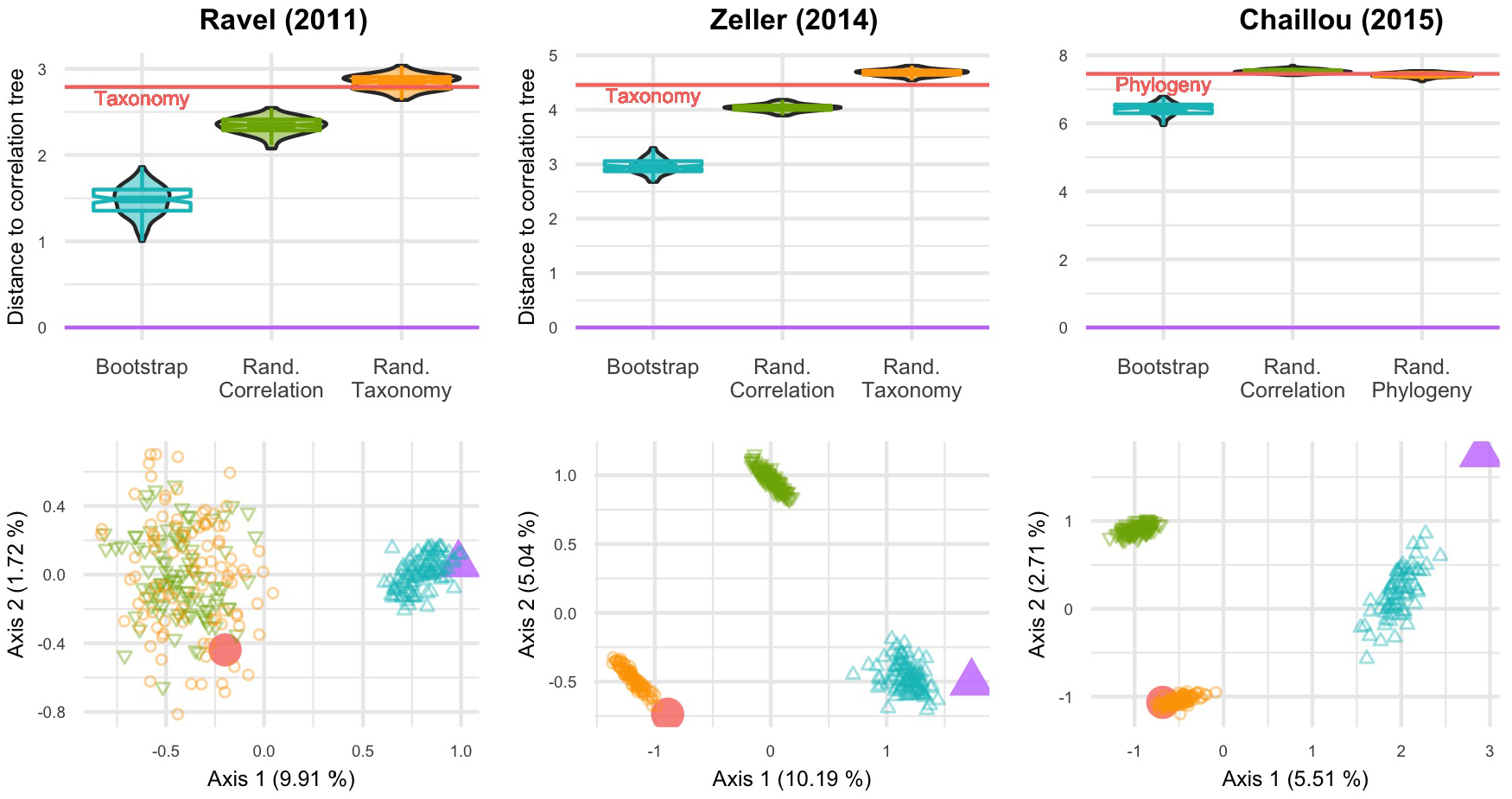
BHV distances between various trees for three datasets: Ravel (left), Zeller (center) and Chaillou (right). Top row: violinplots and notched boxplots of distances to the correlation tree. The distance between taxonomy (or phylogeny) and correlation is indicated by the red line. Bottom row: PCoA projection of all distances on the principal plane. The correlation tree is in purple (Δ), taxonomy (or phylogeny) in red (○), boostraped trees in blue, random correlation trees and random taxonomies (or phylogenies) in green and orange respectively.

Similarly, the PCoA results (Fig. 3, bottom row) highlight two or three tree islands (Jombart et al., 2017): one for the correlation tree and its bootstrap replicates, one for the taxonomy and its randomized replicates and the final one for randomized correlation trees. All random trees can belong to the same island, as seen in the Ravel dataset. The first axis of PCoA represents 5 to 10% of the explained variance and systematically separates the taxonomy from the correlation tree. Moreover, the taxonomy is neither in the bootstrap confidence region of the correlation tree, nor closer to it than a randomized tree.

The only exception is the Chlamydiae dataset, where the phylogeny is within the confidence region of the correlation (Sup. Fig. S1). Note however that this dataset is very small (26 samples) and has many taxa with low abundances, resulting in an extremely large confidence region for the correlation tree. It is also the only one that covers environments ranging from stool to soil and freshwater and thus, for which ecological niche and taxonomy may overlap (Philippot et al., 2010).

In light of these results, we find that the phylogeny is different from the correlation tree, especially when focusing on a single biome. In other words, taxa with similar abundance profiles are not clustered in the phylogeny and the phylogeny may therefore not be a good proxy to find groups of diffentially abundant taxa.

Similar results are observed when using RF distance instead of BHV distance (Sup. Fig. S2).

### 3.2 Pros & Cons of the Different Trees

Athough phylogeny (resp. taxonomy) are evolutionary (resp. ecologically) meaningful and increasingly available, they do not capture similarities between taxa in terms of abundance profiles. For example, if abundances are driven by a phenotype regulated by a mobile element (*e.g.* an antibiotic resistance gene), evolutionary and ecological histories are not informative. Furthermore, when performing differential abundance analyses with genes (metatranscriptomics) or metagenomics-based taxa such as MSPs and metagenome-assembled genomes, many of which are poorly annotated, neither a taxonomy nor a phylogeny is available.

In contrast, the correlation tree is constructed from the abundance data and can thus always be used. By its very definition, it clusters taxa with similar abundance profiles. Unfortunately, it suffers from limitations of its own. First, it is estimated from the data and thus sufficient data should be available to build a robust correlation tree. Second, since the same data are used to build the correlation tree and to test differential abundance, some care should be taken not to overfit the data. For example, permutation-based tests are valid because the group labels are not used during the tree construction and are thus independent of the hierarchical structure (Goeman and Finos, 2012) but other tests should be used with caution.

### 3.3 Simulation Study

#### 3.3.1 Non-Parametric Simulations

Note first that *z*-smoothing numerically fails and does not produce any results in an average 4% of the simulations (ranging from 2% for the randomized correlation tree to 8% for the correlation trees). Second, the hyperparameters *k* and *ρ* controlling the level of smoothing are often very far from 1 (below and above, respectively) resulting in little to no smoothing. Fig. 4 shows the impact of smoothing on *z*-scores: in more than half of the simulations, the *z*-scores were shifted by less than 10^−2^ units in either direction. Among the different topologies tested, the phenomenum was the strongest for the correlation trees: the *z*-scores were shifted by more than 10^−2^ units in less than 5% of the simulations.

**Figure 4.**
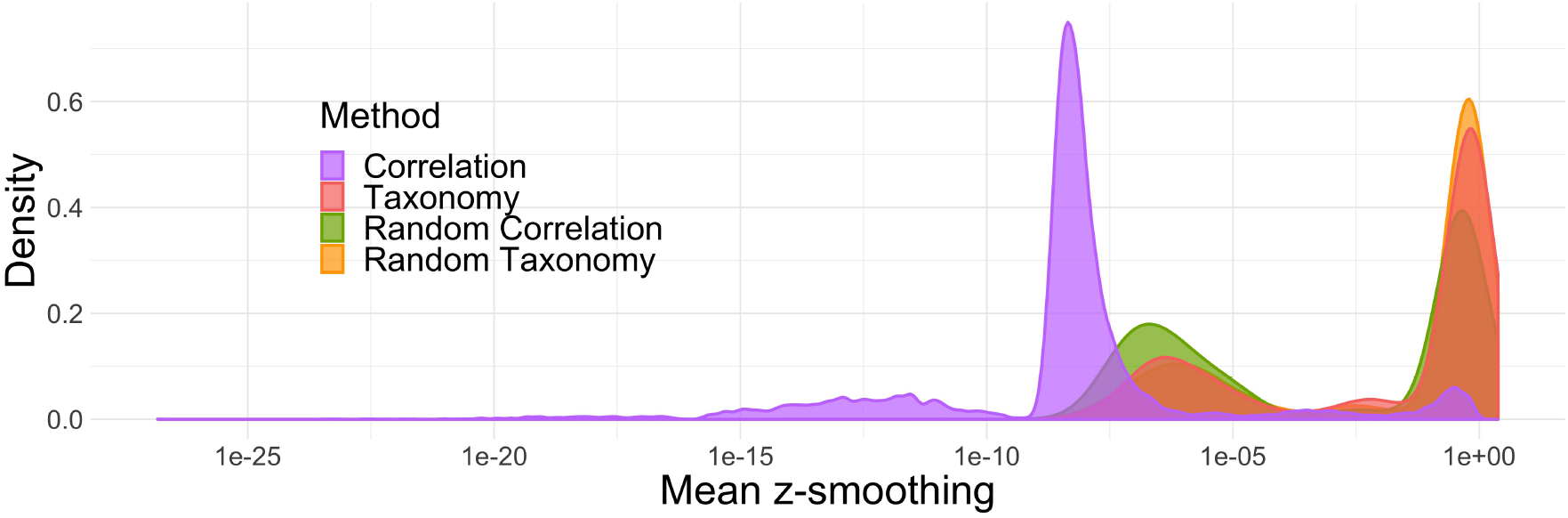
Average absolute difference between *z*-scores before and after smoothing. In most simulations, smoothing only marginally changes the results.

Concerning FDR control, the standard BH procedure was the only one that achieved a nominal FDR rate below 5% across different fold changes and proportions of null hypothesis (Fig. 5, bottom row). All other procedures exceeded the target rate, reaching nominal rates of up to 7%, when the number of null hypothesis grew beyond 90%.

**Figure 5.**
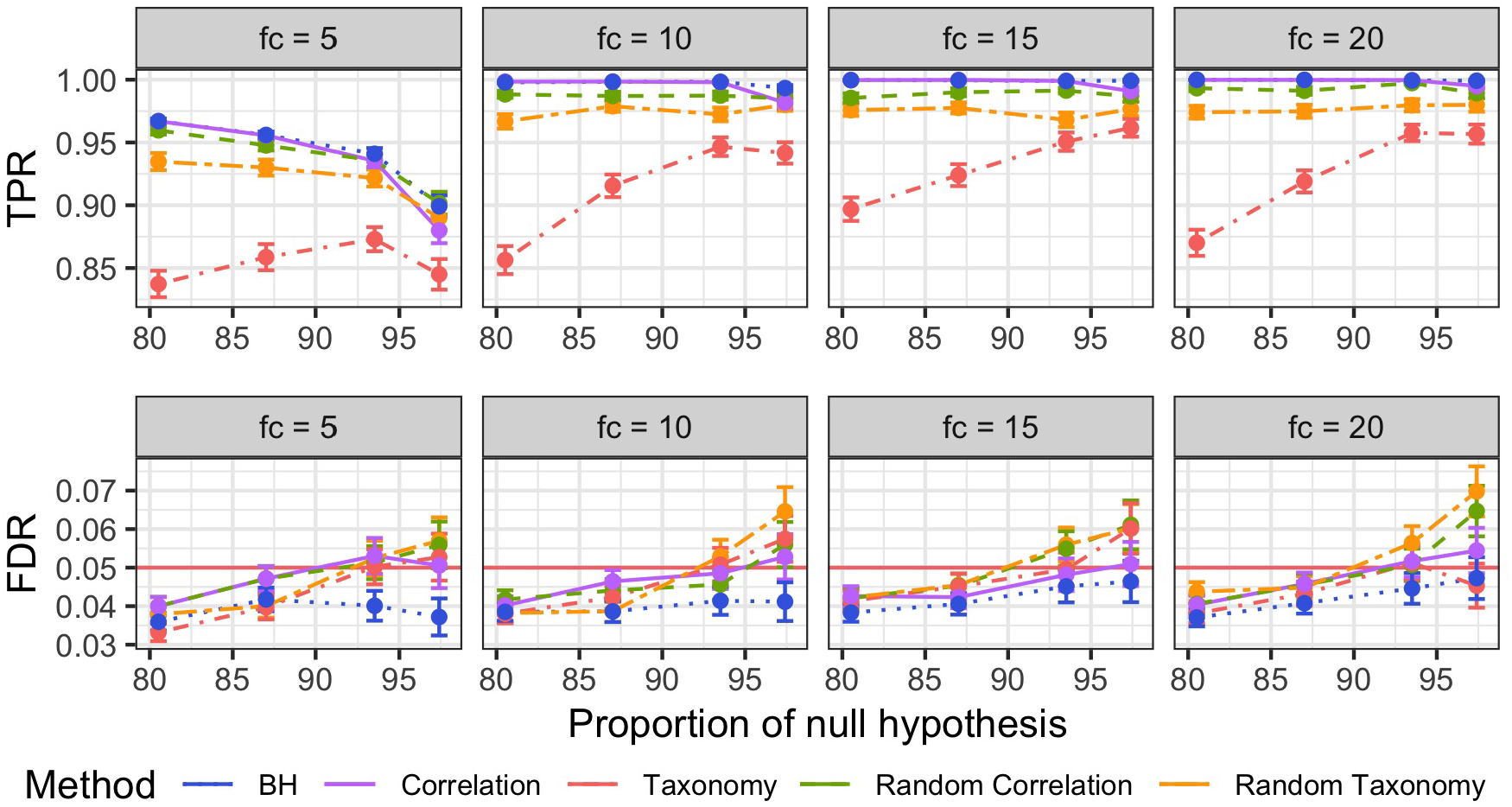
Mean and Squared Error of the Mean (SEM) of the true positive rates (TPR, top) and FDR (bottom) per different fold changes (facets) for non-parametric simulations. The different FDR control procedures are color-coded. Mean and SEM are computed over 600 replicates.

BH was similarly the most powerful method across all fold changes and proportions of null hypothesis (Fig. 5, top row), with correlation tree and randomized correlation trees coming close second and third. BH, correlation tree and randomized trees outperformed the taxonomy in all settings, resulting in TPR increase of up to 0.15.

The quasi-equivalence between BH and correlation tree is not surprising given the absence of smoothing when using the correlation tree. The comparatively bad result of the taxonomy is also expected from our simulation settings as the taxonomy is independent from simulated differential abundance. Forcing the discoveries to be close in the tree therefore introduces a systematic bias and results in a loss of power, especially for differential taxa that are isolated, and an increase in false discoveries, especially for non-differential taxa that are close to differential ones.

The better results of *a priori* uninformative random trees compared to the taxonomy were however more surprising, especially in light of the similar levels of smoothing for all those trees. It turned out that the random trees were, on average, closer to the correct correlation structure of differential taxa than the taxonomy and therefore had a lesser negative impact on the detection power.

It is clear from these results that using a tree reflecting the true data structure, such as the correlation tree, does not increase the number of discoveries but does not degrade the perforance of the method either. In contrast, using a wrong structure degrades the detection power from only slightly at best (for random trees) to quite a lot (taxonomy).

#### 3.3.2 Parametric Simulations

Parametric simulations showed exactly the same patterns as non-parametric ones. *Z*-scores smoothing was limited in most replicates and almost always null when using the correlation tree (Supp. Fig. S3). BH was the only procedure with a nominal FDR below the target rate of 5% in all settings and all trees led to nominal above the threshold when the proportion of differential taxa was low (Supp. Fig. S4, bottom row). Finally, BH had the highest TPR among all methods (Supp. Fig. S4, top row).

The results differed from the non-parametric ones in one important aspect: all methods had low TPR, below 0.15, whereas they achieve TPR higher than 0.85 in the non-parametric setting. This difference is mainly due to the parametric simulation scheme, reused from Xiao et al. (2017): differential taxa are not pre-filtered based on their prevalence and can thus have a very high proportion of zeros in the worst case. Multiplication by a fold-change, no matter how high, leaves those zeroes and their corrresponding ranks unchanged. This in turn strongly degrades the ability of the rank-based Wilcoxon test, to find differences between groups among those taxa.

### 3.4 Analysis of Real Datasets

#### 3.4.1 Reanalysis of Chlamydiae dataset

The Chlamydiae dataset consists of 26 samples distributed over 9 very different environments (feces, freshwater, human skin, sea, …). Differential abundance of the OTUs across the environment was tested using the same parameters as in the original article (hFDR on the phylogeny, *α* = 0.1). The test identified 8 differential OTUs with a global *a posteriori* FDR of *α*′ = 0.32. Substituting the correlation tree to the phylogeny in this analysis led to the detection of 3 additional OTUs, at a comparable global FDR of *α*′ = 0.324.

Abundance boxplots of these three additional OTUs (Fig. 6, insets **E** and **F**) show that these OTUs are much more abundant in soil samples and almost specific to that environment, validating their differentially abundant status. In that example, the correlation tree reflected the structure of the data better than the phylogeny and increases the power at no cost to the nominal FDR.

**Figure 6.**
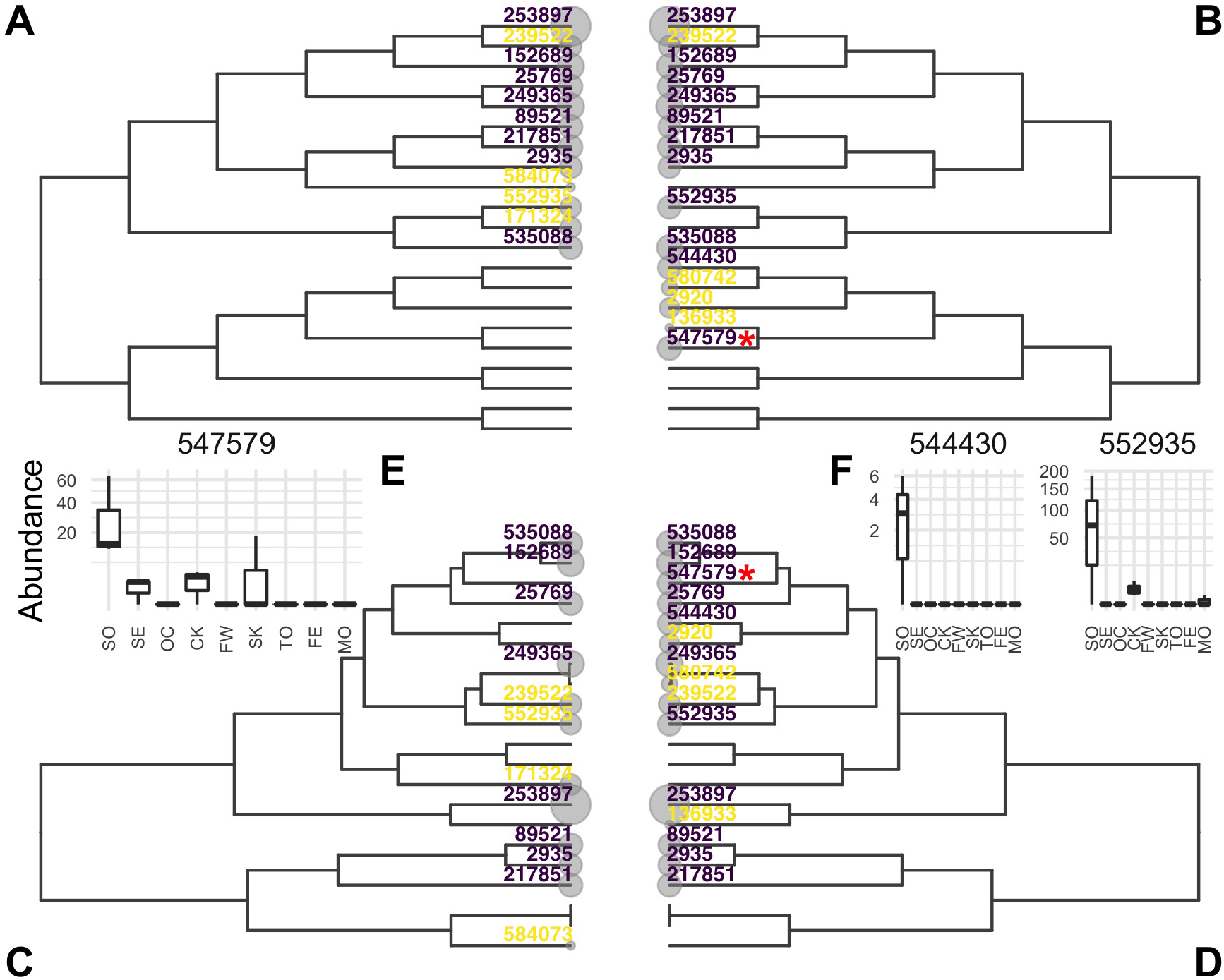
**A**-**D**: Evidences of OTUs estimated by hFDR with phylogeny (**A** and **C**) or correlation tree (**B** and **D**) represented on phylogeny (**A** and **B**) or correlation tree (**C** and **D**). OTUs detected as differential are colored in purple, those tested but not detected as differential in yellow. **E**-**F**: Abundances of OTUs detected only by the correlation tree in different environments. OTU 547579 in **E** is hightlighted with a red star in **B** and **D**. Environment are abbreviated as SO: soil, SE: sediment, OC: ocean, CK: creek, FW: fresh water, SK: skin, TO: tongue, FE: feces, MO: mock.

Fig. 6 shows the location of evidences (*e* = − log_10_(*p*)) and differential OTUs on both the phylogeny and correlation trees. OTU 547579, highlighted with a red star, is one the three additional OTUs. It was not tested with the phylogeny because it is the only differential taxa in its clade (panel **B**) and its top-most ancestor was not rejected. In contrast, it belongs in the correlation tree to a group of soil-specific taxa and the hierarchical procedures sequentially rejected all its ancestors so that it was also tested and rejected.

With this top-down approach, the correlation tree is a better candidate hierarchy than the phylogeny. Indeed, the signals of differential OTUs can be averaged out with noise and/or conflicting signal in the phylogeny, they are pooled together in the correlation tree. This makes it easier to reject high level internal nodes and descend the tree toward differential OTUs.

It should be noted however that the *a posteriori* global FDR is quite high at 0.324. Using the standard BH with a FDR of 0.324 results in 4 new discoveries, for a total of 15. hFDR, with either the correlation or the phylogeny, does not outperform the classical BH procedure. This discrepancy might be explained by the global FDR computation used in hFDR which controls the FDR in the worst case scenario. The actual global FDR could be much lower than this pessimistic bound.

#### 3.4.2 Analysis of Chaillou dataset

The Chaillou dataset consists of 64 samples uniformly distributed across 8 food types (ground veal, ground beef, poultry sausages, sliced bacon, shrimps, cod fillet, salmon fillet, smoked salmon). Differential abundances of OTUs from the Bacteroidetes phylum (97 OTUs) across food types was tested with hFDR procedure (*α* = 0.01, both phylogeny and correlation tree). The test had a global *a posteriori* FDR of 0.04 for both the phylogeny and the correlation tree and detected 28 differential OTUs with the phylogeny and 34 with the correlation tree. Similarly, with a 0.04 FDR level, vanilla BH leads to 55 discoveries.

Unlike the Chlamydiae dataset, only 22 OTUs were detected by both methods. Careful examinations of those 22 show that each of them (i) is missing, or below the detection level, in at least one of the 8 food type of the studies whereas and (ii) has high prevalence (≥0.75%) and abundance in at least one other food type. We can thus classify those 22 as true positives rather than false discoveries.

The abundance profiles of the 18 OTUs found only by the correlation tree (hereafter cor-OTUs) or the phylogeny (phy-OTUs) (Sup. Fig. S5) show marked differences across the the 8 food types, validating their differential status. As was the case in the Chlamydiae dataset, cor-OTUs are often isolated in the phylogeny (Sup. Fig. S6) and thus not even tested during the hierarchical procedure as they are averaged with low-signal taxa.

In contrast, phy-OTUs are often close to detected taxa in the correlationtree but not detected because of the *F*-test implemented in StructSSI. For example, the three phy-OTUs 0656, 1495 and 0241 belong to a cluster of five shrimp-specific OTUs but the two others (0516 and 0519) have some outlier counts and comparatively higher counts that the three phy-OTUs (Sup. Fig. S7, right). Aggregation at internal nodes leads to high variance which decreases the significance of the *F*-test: *p*-values at the internal nodes do not pass the threshold and the leaves are not tested. Replacing the *F*-test with the Kruskal-Wallis test, which is more robust to outliers, led to the detection of all OTUs (Sup. Fig. S7, left).

#### 3.4.3 Analysis of genera in Zeller dataset

The Zeller dataset consists of gut microbiomes from 199 subjects that are healthy (*n* = 66), suffer from adenomas (*n* = 42) or from colorectal cancer (*n* = 91). Differential abundances of genera across medical conditions was tested with *z*-score smoothing, using several tree (no tree or standard BH, taxonomy, correlation tree, randomized correlation tree and randomized taxonomy) and several FDR threshold levels.

Fig. 7 (left panel) shows the number of genera detected by each tree at each threshold. While the correlation tree detects the most taxa and BH the least at almost all threshold values, the differences between all trees are very small (one or two taxa only). In particular, at *α* = 0.05, all methods detected either 14 or 16 genera.

**Figure 7.**
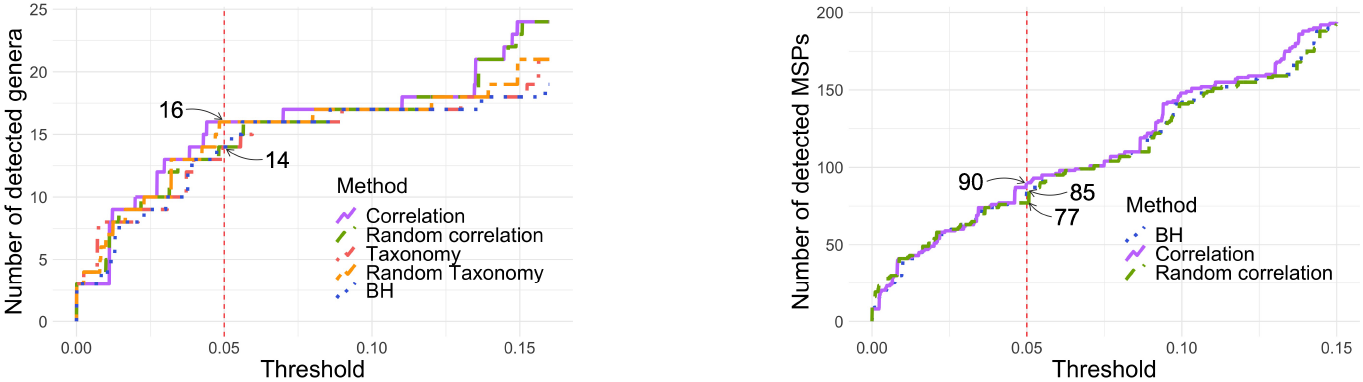
Number of detected genera (left) or MSPs (right) according to the *p*-value threshold. Left: with *α* = 0.05, 14 genera are detected with taxonomy, random correlation tree and BH while 16 species are detected with correlation tree and random taxonomy. Right: with *α* = 0.05, 85 MSPs are detected by BH and 90 by correlation tree.

In this example, the algorithm estimated *ρ* > 40 for the random trees and *k* < 10^−7^ for the correlation tree, effectively resulting in no smoothing of the *z*-scores. The corresponding values are *ρ* = 0.26 and *k* = 0.37 for the taxonomy. The *z*-scores were thus smoothed to a higher extent but this had almost no impact on the number of detected genera.

#### 3.4.4 Analysis of MSPs in Zeller dataset

Repeating the same analysis at the MSP, rather than genus, level gave similar results. Among the 878 MSP and using *α* = 0.05, 234 were detected without correction, 90 with the correlation tree, 85 with standard BH and 77 with a random tree. Neither the taxonomy nor the phylogeny were available for the MSP and they were therefore not compared to the other methods.

In that example *k* = 1.3 × 10^−7^ and the tree has almost no impact on the *z*-scores and the corrected *p*-values (Sup. Fig. S8, bottom row). The 5 additional taxa detected with the correlation tree are indeed not clustered with other detect taxa and have BH-corrected *p*-values between 0.0505 and 0.0540 (Sup. Fig. S8, left row). The main differences between the two procedures does not lie in the use of a hierarchical structure rather than in the way corrected *p*-values are computed: using permutations for the correlation and analytic formula for BH. It coincides with previous findings that permutation-based FDR control improves detection of differentially abundant taxa (Jiang et al., 2017).

## 4 Conclusion and perspectives

In this work, we investigated the relevance of incorporating *a priori* information in the form of a phylogenetic tree in microbiome differential abundance studies. Doing so was reported to increase the detection rate in recent work (Xiao et al., 2017; Sankaran and Holmes, 2014).

The rationale rests upon the assumption that evolutionary similarity reflects phenotypic similarity. Taxa from the same clade should therefore be more likely to be simultaneously associated to a given outcome than distantly related taxa. Although this assumption sounds natural and supported by evidence for high level taxa such as phylum (Philippot et al., 2010), there are also many arguments against it for low level taxa such as species and strains. Previous work (Harris et al., 2014) even showed some degree of equivalence between species in the gut, *i.e.* species within the same ecological guild could replace each other during the assembly process.

We considered here whether the phylogeny and taxonomy were good *a priori* trees to capture the structure of the abundance data, as captured by the correlation tree. In all the environments we studied, we found that the taxonomy and/or the phylogeny were significantly different from the correlation tree. Taxa with very similar abundance profiles could be widely spread in the phylogeny and vice-versa. The phylogeny was on average no closer to the correlation tree than a random tree, and thus not a good proxy of the abundance data structure.

We further studied the impact of tree misspecification on two recently published tree-based testing procedures, *z*-score smoothing (Xiao et al., 2017) and hFDR top-down rejection (Yekutieli, 2008).

Concerning *z*-score smoothing, we showed on synthetic data that substituting the correlation tree to the phylogeny increased the detection rate. Quite surprisingly, replacing the phylogeny with a random tree also increased the detection rate (Fig. 5), questioning the use of the phylogeny in the first place. The results were even more disappointing on real datasets where all trees led to similar detection rates and none of them significantly outperformed standard BH (Fig. 7). In the Zeller MSP dataset, the differences between procedures were limited (Sup. Fig. S7) and stemmed mostly from the way p-values were computed: *i.e.* using permutations for *z*-score smoothing and closed formula for BH. Overall, using phylogenetic information to smooth *z*-scores degrades the detection rate (at worst) or leaves it unchanged (at best).

Top-down rejection (hFDR) gave more interesting results. Replacing the phylogeny or taxonomy with the correlation tree increased the detection rate, while preserving the global *a posteriori* FDR. In general, taxa detected with the correlation tree but not with the phylogeny belonged to clades of mostly non-differential taxa in the phylogeny (Fig. 6). Their signal was thus averaged with noise and they discarded early-on in the hierarchical procedure. In contrast, they were salvaged on the correlation tree as they belonged clades of taxa with similar abundance profiles. Unfortunately, hFDR suffers from two limitations. First, it has a lower detection rate than standard BH at the same global FDR level. This is likely a side effect of the definition of the global FDR in hFDR, *i.e.* FDR in the absolute worst case scenario. Second, the current implementation of hFDR in `StructSSI` is limited to *F*-test, which are ill-suited to highly non-gaussian microbiome data.

Our conclusions are two-fold. First, the phylogeny does not capture the structure of the abundance data and should be replaced by a better hierarchical structure such as the correlation tree. Second, hierarchical methods in their current state do a poor job of leveraging the hierarchical information to increase the detection rates. Until better hierarchical methods are available (*e.g.* hFDR with support for more complex tests), we recommend sticking to the time-tested BH procedure for differential abundance analysis.

## Author Contributions

MM, CA and JP designed and directed the study. AB, MM, CA and JP wrote the manuscript. AB created the synthetic datasets. AB performed all the analyses with substantial input from MM, CA and JP. All authors discussed the results and commented on the manuscript.

