## Supplemental Material for "Incorporating phylogenetic information in microbiome abundance studies has no effect on detection power and FDR control"

Antoine Bichat [1,2], Jonathan Plassais [2],
Christophe Ambroise [1] and Mahendra Mariadassou [3,*]
[1]LaMME, Université d'Évry val d'Essonne, 91000 Évry, France
[2]Enterome, 94-96 Avenue Ledru Rollin, 75011 Paris, France
[3]MaIAGE, INRA, Université Paris-Saclay, 78350, Jouy-en-Josas, France

### 1 Reproducibility

A R-package was been made to provide helpful functions to support the analysis performed in this work. It is available on the GitHub repository: `https://github.com/abichat/correlationtree`.

All codes for analysis and figures are also available on GitHub: `https://github.com/abichat/correlationtree_analysis`. Some results or figures might be slightly different due to a different random number seed choice or a different number of simulations.

### 2 Supplementary Tables and Figures

#### 2.1 Figures

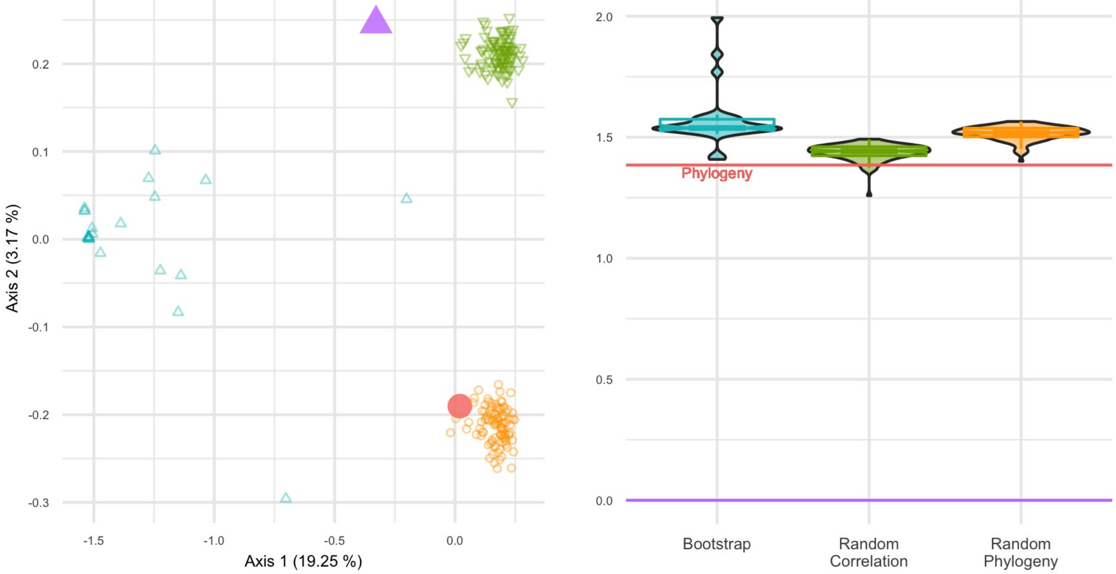


Figure 1 BHV distances on forest of trees generated on the Chlamydiae dataset. Left: PCoA of pairwise distances. Right: distance to the correlation tree. Unlike other datasets studied in the analysis, bootstrap replicates of the correlation tree are quite far from the original one, and no closer than the phylogeny.

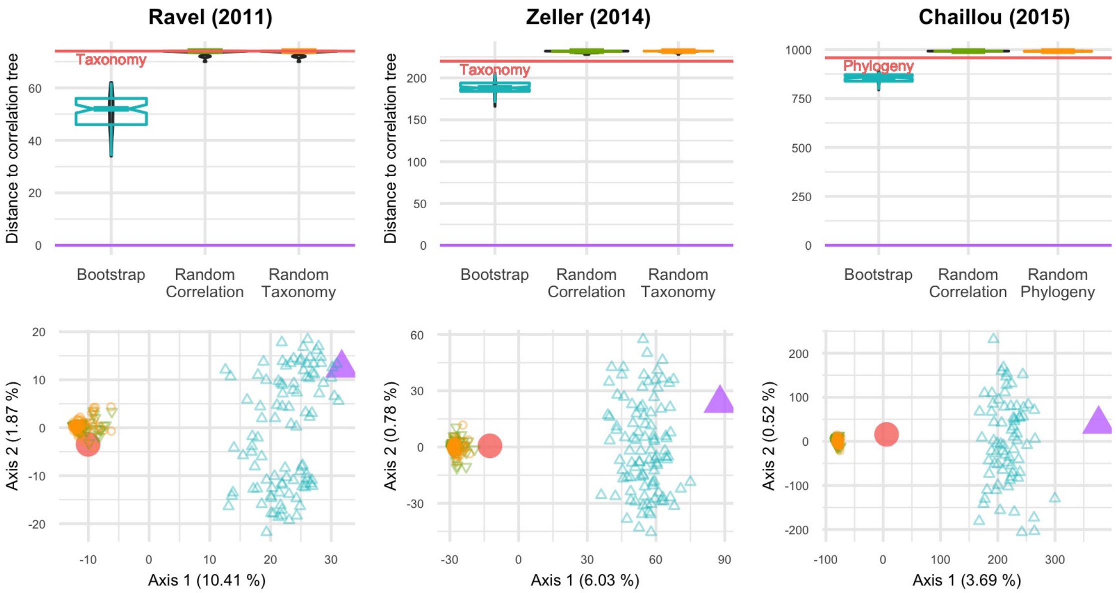


Figure 2 RF distances between the correlation tree and various other trees for three datasets: Ravel (left), Zeller (center) and Chaillou (right). Top row: violinplots and notched boxplots of distances to the correlation tree. The distance between taxonomy (or phylogeny, depending on the dataset) and correlation corresponds to the red line. Bottom row: PCoA projection of all distances on the principal plane. The correlation tree is in purple (△), taxonomy (or phylogeny) in red (○), boostraped trees in blue, random correlation trees and random taxonomies (or phylogenies) in green and orange respectively. The first axis of the PCoA always separates the taxonomy / phylogeny from the correlation tree.

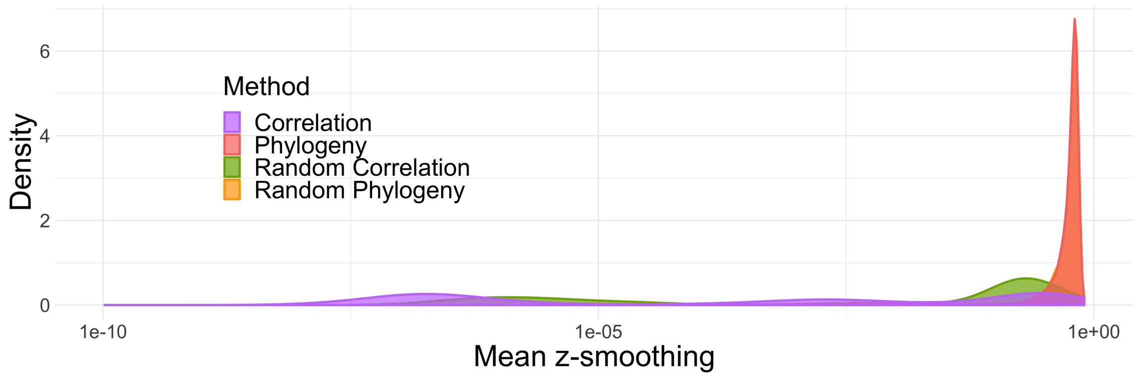


Figure 3 Average absolute difference between $z$-scores before and after smoothing for parametric simulations. Phylogeny and random phylogenies densities perfectly overlap. Int most simulations, the $z$-scores are barely changed by the smoothing procedures.

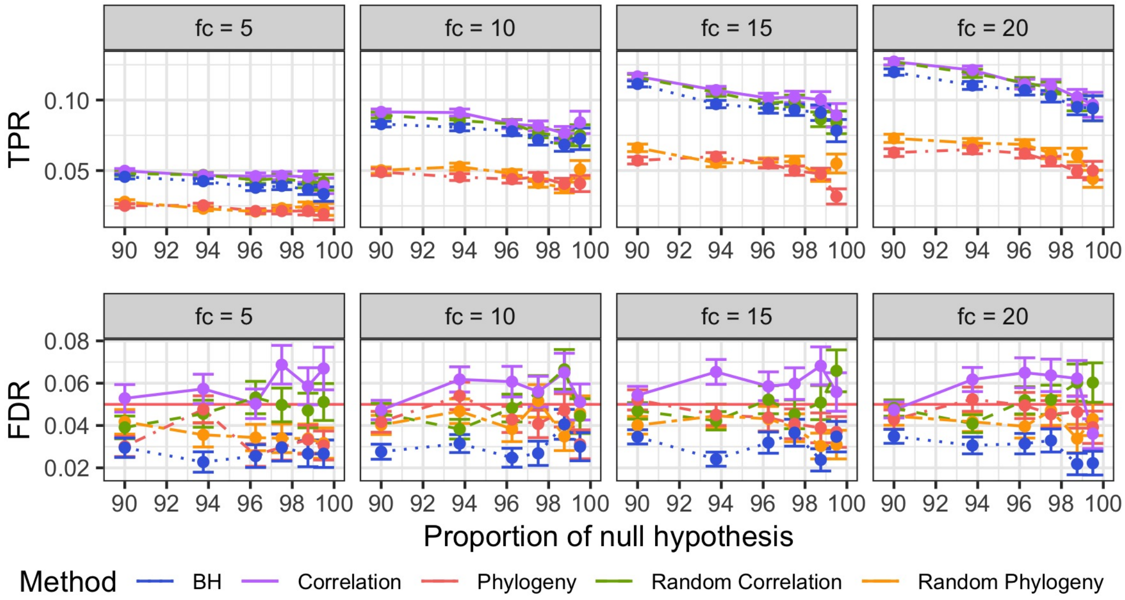


Figure 4 Mean and Squared Error of the Mean (SEM) of the true positive rates (TPR, top) and FDR (bottom) per different fold changes (facets) for parametric simulations. The different FDR control procedures are color-coded. Mean and SEM are computed over 600 replicates. BH and the correlation tree always outperform the phylogeny but BH is the only one to achieve a nominal FDR below 0.05.

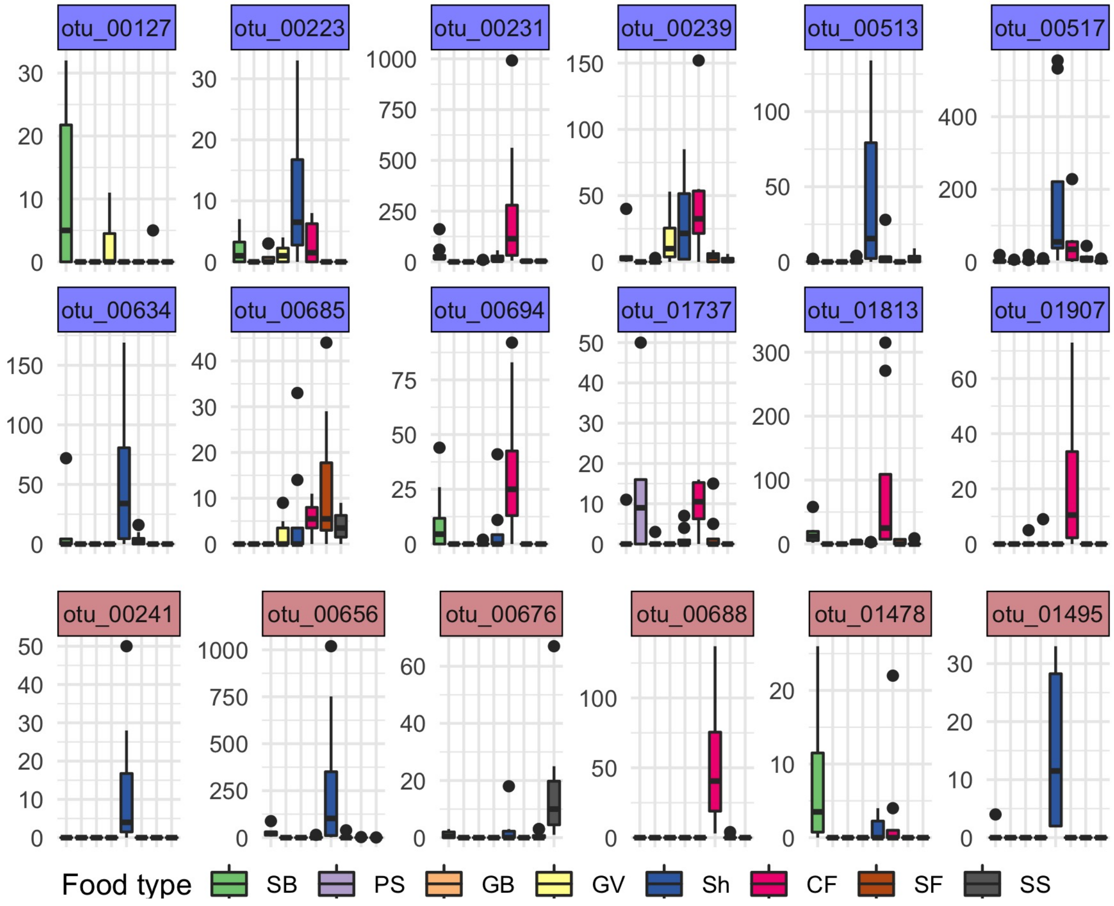


Figure 5 Abundances for OTUs detected only by the correlation tree (topmost rows, blue strip background) or by phylogeny (bottom row, red strip background) in the Chaillou dataset. Food types are abbreviated as SB: sliced bacon, PS: poultry sausage, GB: ground beef, GV: ground veal, Sh: shrimp, CF: cod fillet, SF: salmon fillet, SS: smoked salmon. All 18 OTUs have very different abundances across food types.

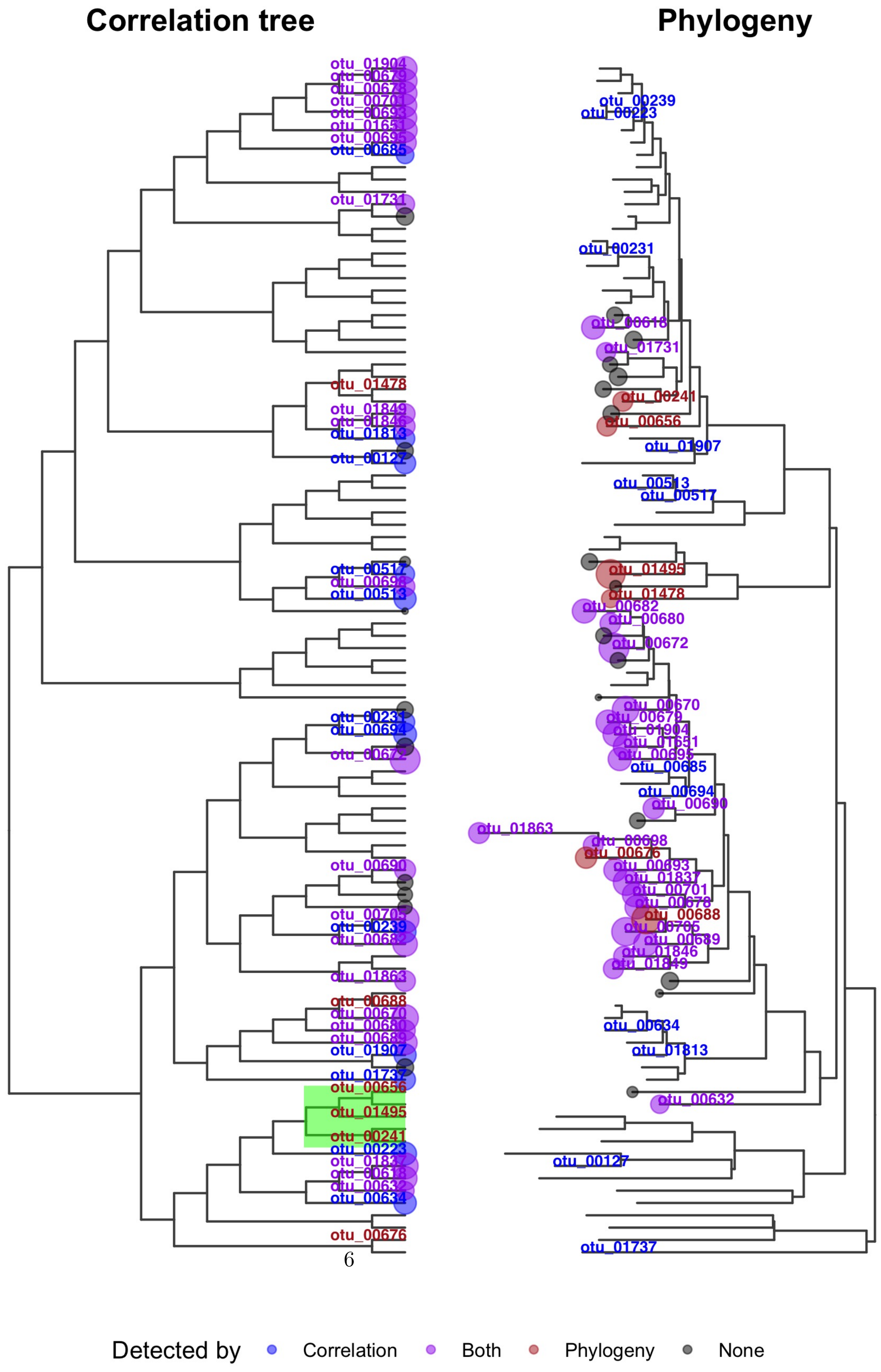


Figure 6 Position of the detected OTUs on both the correlation tree (left) and the phylogeny (right). Point sizes are proportional to evidences (-$log_{10}$($p$-

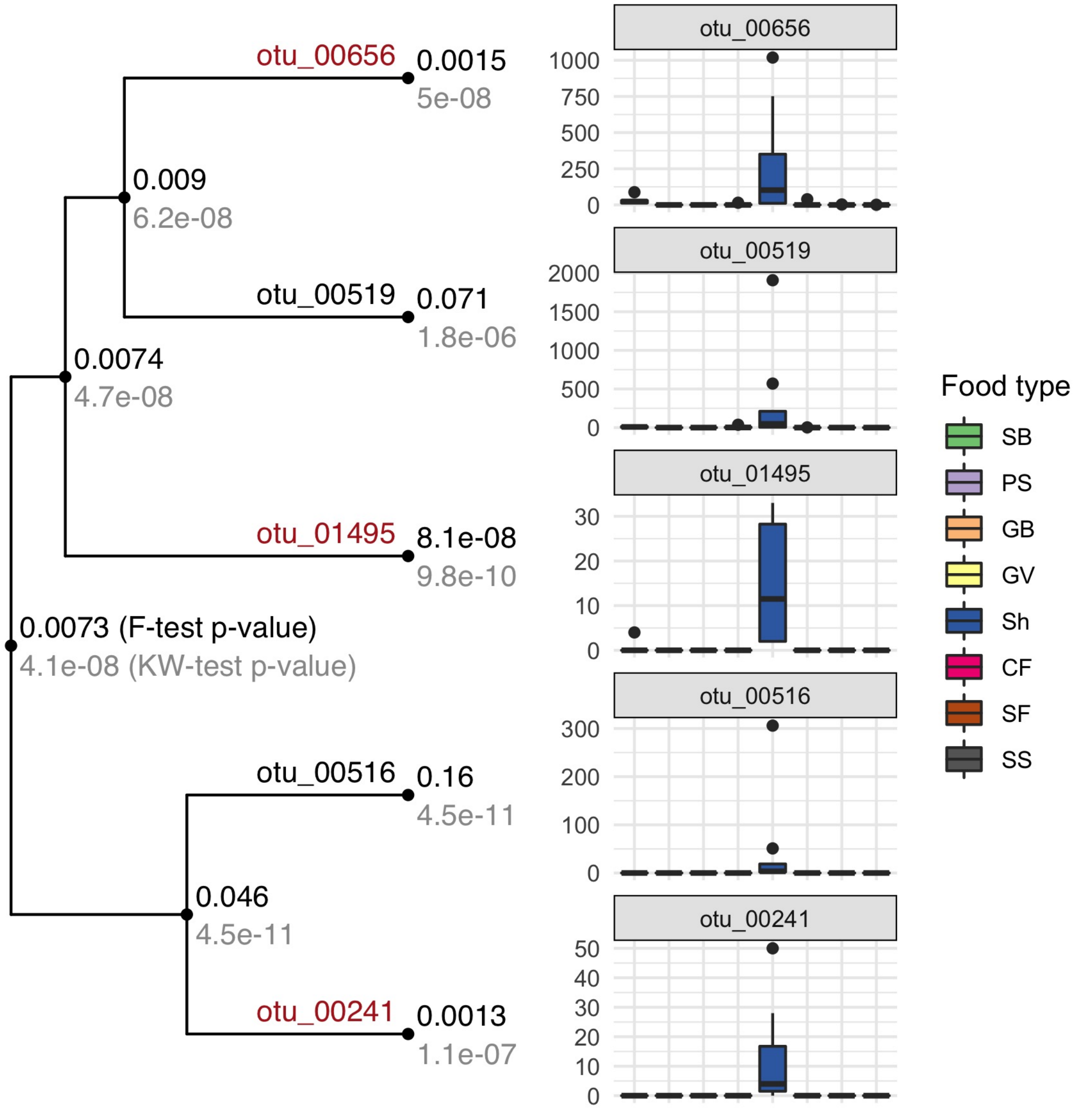


Figure 7 Focus on the green clade highlighted in Sup. Fig. S6. Left: Tree topology and $p$-values from a Fisher test (top) and a Kruskal-Wallis test (bottom) at each tip and each internal node of the correlation tree. Abundances of internal nodes were computed by summing the abundances of all leaves in their subtree. Right: Abundance profile of each OTU across food types. Food types are abbreviated as SB: sliced bacon, PS: poultry sausage, GB: ground beef, GV: ground veal, Sh: shrimp, CF: cod fillet, SF: salmon fillet, SS: smoked salmon. $F$-test are not significant due to (i) outlier counts in many abundance profiles and (ii) the obvious differences in variance across food types.

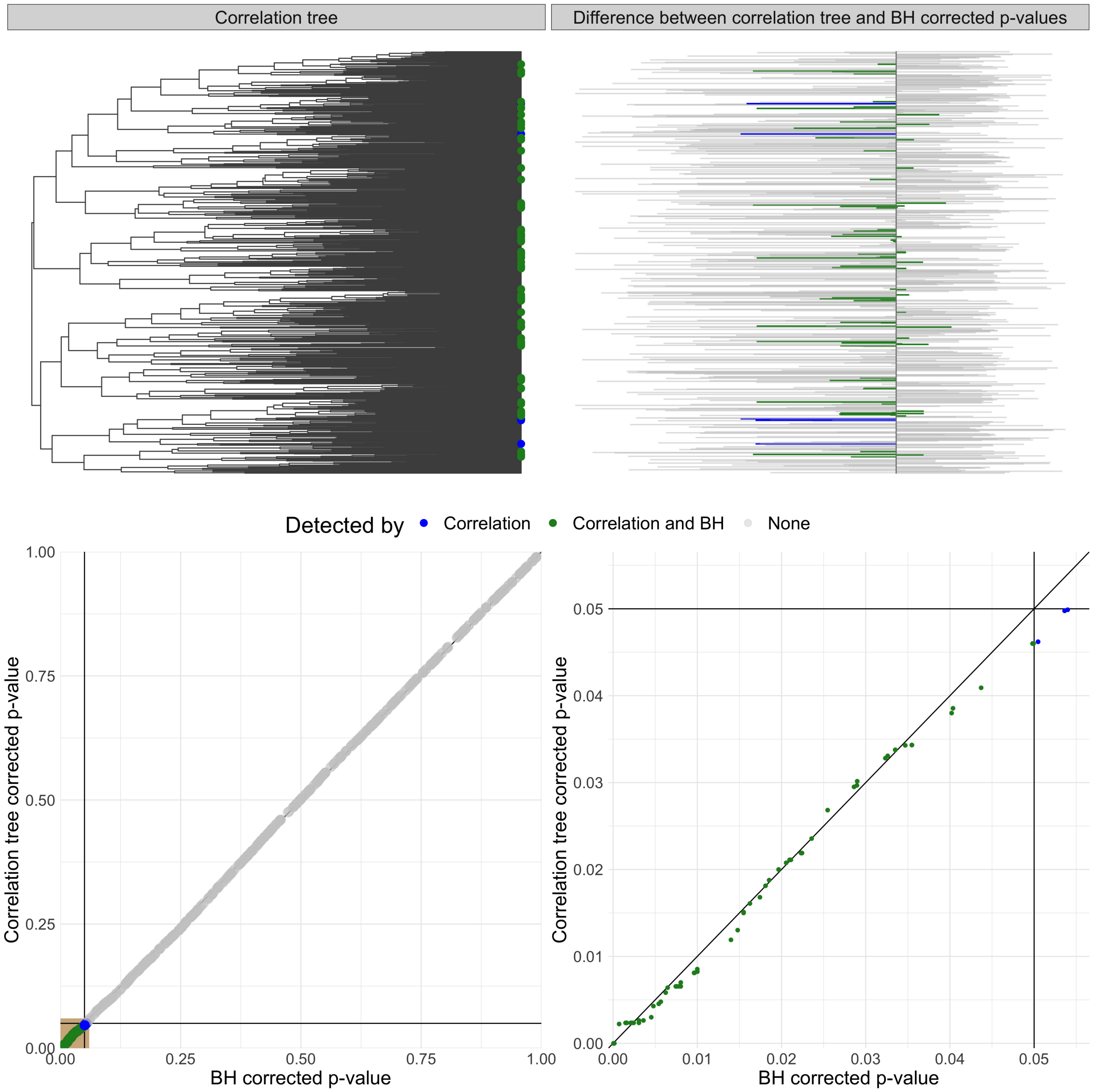


Figure 8 MSP from Zeller detected only by the correlation tree (blue) or also by the BH correction (green). Top left: Detected OTUs and their location on the tre. Top right: difference beetween $p$-values corrected by the correlation tree and by BH. Bottom left: $p$-values adjusted with the correlation tree against those adjusted with the BH correction. Bottom right: zoom on significant $p$-values (tan box). $z$-scores are virtually uncorrected ($k = 1.3 \times 10^{-7}$) leading to almost identical $p$-values. The difference between the two sets of $p$-values lies in the use of different correction methods: permutation-based for the correlation tree, formula-based for BH.
